# A temporal parcellation of the sensory-evoked responses during the rubber hand illusion reveals manipulation- and illusion-specific correlates

**DOI:** 10.1101/2021.01.15.426770

**Authors:** Placido Sciortino, Christoph Kayser

## Abstract

The neurophysiological processes reflecting body illusions such as the rubber hand remain debated. In particular, previous studies investigating neural responses evoked by the illusion-inducing stimuli provide diverging reports as to when these responses reflect the illusory state. To resolve these controversies in previous work we applied multivariate (cross-) classification to EEG responses obtained during the rubber hand illusion and multiple control conditions in human participants. These controls were designed to test for markers of the illusory state that generalize across the spatial arrangements of limbs or the specific nature of the control condition (rubber hand or participant’s real hand) - hence which are independent of the precise experimental conditions used as contrast for the illusion. This revealed a parcellation of evoked responses into a temporal sequence of events that each differentiate the illusion and control conditions along distinct dimensions. Importantly, around 130-150 ms following stimulus onset the neurophysiological signals reliably differentiated the illusory-state from all non-illusion epochs. This neurophysiological signature was not correlated with changes in skin conductance accompanying the illusion, suggesting that neurophysiological and bodily signals reflect distinct illusion-related processes.

## Introduction

The neurophysiological processes underlying multisensory body perception and the sense of body ownership are often studied using illusions such as the rubber hand (Blanke, 2012; Botvinick and Cohen, 1998; Longo and Haggard, 2012; Riemer et al., 2019). There, participants watch an artificial hand being stimulated in synchrony with their own occluded hand. This synchronous sensory stimulation of their own and the rubber hand results in the illusory experience of the rubber hand becoming embodied (Blanke, 2012; Tsakiris and Haggard, 2005). Despite many behavioural studies demonstrating the robustness of the illusion to variations in the inducing setup, such as the relative spatial arrangement of hands or the use of real-world or artificial objects like avatars (Bertamini and O’Sullivan, 2014; Kalckert and Ehrsson, 2012; Tsakiris and Haggard, 2005), the precise neurophysiological correlates of the rubber hand illusion (RHI from now) remain controversial.

Functional imaging studies have linked a broader network of parietal, premotor and frontal regions to body illusions (Bekrater-Bodmann et al., 2014; Ehrsson, 2004; Petkova et al., 2011). However, previous electrophysiological studies have disagreed on the relevant illusion-specific activations. For example, several studies have investigated the sensory-evoked potentials produced by the repetitive visuo-tactile stimulation, and asked when and how these evoked responses are affected by the illusory state - the subjective state of the individuals when they experience the illusion that is absent in the control conditions. One study reported illusion-correlates at early (∼50ms) latencies relative to the stimulation events and advocated for a low-level origin of the illusion related processes in early somatosensory cortex (Zeller et al., 2015). Support for such a low-level origin comes also from studies on the general enhancement of somatosensory signals by simultaneous visual stimuli (Cardini et al., 2012, 2011). However, other studies advocated for illusion-correlates only at latencies of around 120-200ms, possibly related to the detection of mismatching sensory signals in parietal cortex (Aspell et al., 2012; Press et al., 2008). A third group of studies reported illusion-related activity only at latencies of around 300-400ms and suggested high-level cognitive processes as respective sources (Peled et al., 2003; Rao and Kayser, 2017). And finally, a recent study on self- and agent-generated tactile inputs reported no differences in evoked responses pertaining to the RHI (Pyasik et al., 2021). Hence, it remains unclear when following stimulus onset the illusion-specific activations occur.

One reason for such conflicting results may be the different procedures used to induce the illusion in previous work. For example, the illusion has been induced when the real hand is besides (Bekrater-Bodmann et al., 2014; Kammers et al., 2011; Lloyd, 2007; Rao and Kayser, 2017; Riemer et al., 2015) or below the rubber hand (Preston, 2013; Rohde et al., 2011; Zeller et al., 2015). Yet, few studies asked which neurophysiological correlates generalize across distinct procedures to induce the illusion. A second reason can be the stimulation technique itself: while manual stimulation may provide a more realistic sensation, computer-controlled devices can also induce the illusion (Kanayama et al., 2021; Rao and Kayser, 2017; Suzuki et al., 2013) and allow for a more precise alignment of rapid neurophysiological processes to the somatosensory stimulus. Third, previous studies investigating evoked neurophysiological responses during the RHI made the implicit assumption that illusion-related activations have the very same spatial configuration across participants. However, individual variability in brain morphology may invalidate this assumption (Eichert et al., 2020; Li et al., 2019; Van Horn et al., 2008) calling for the use of analytical methods relaxing this assumption, such as multivariate classification (Parra et al., 2005).

The goal of this study was to determine the (neuro-)physiological correlates of the illusory state during the RHI in EEG-derived evoked responses while overcoming the above-mentioned limitations. To this end we employed a computer-controlled setup to induce the illusion along both the vertical and horizontal planes (Rao and Kayser, 2017). Importantly, to determine which neurophysiological signatures generalize across experimental conditions, we combined two traditional illusion-inducing conditions featuring the synchronous stimulation of real and artificial hands with two control conditions: a rubber hand tilted by 90°, and participant’s real hand. Furthermore, we also introduced a control condition devoted to comparing evoked responses within the experimental illusion configuration but between the period before and after participants reporting the onset of the illusion. This analysis can reveal changes in the neurophysiological processes accompanying the illusion that are independent of changes in the external sensory input. To determine the neurophysiological correlates of the illusory state we used multivariate classification, an approach now routinely exploited to uncover the neurophysiological signatures of sensory and cognitive processes (Cichy et al., 2014; Grootswagers et al., 2018; Guggenmos et al., 2018; Keitel et al., 2020). Importantly, this allowed us to directly probe which neurophysiological processes generalize across the spatial configurations of hands when inducing the illusion or across multiple control conditions using cross-classification.

We complemented the EEG recordings with measurements of skin conductance and questionnaires. Skin conductance has been shown to index autonomous responses that are sensitive to the embodiment of the rubber hand (Armel and Ramachandran, 2003; Critchley et al., 2021; D’Alonzo et al., 2020; Ehrsson et al., 2007). This also allowed us to ask whether the neurophysiological (EEG) correlates of the illusory state are related to those physiological correlates obtained from the skin conductance.

## Materials and methods

### Experimental Conditions

Experiments were performed in a darkened and electrically shielded room (Ebox, Desone, Germany). Participants sat on a comfortable chair in front of a one-compartment, open-ended box placed on a two-story wooden platform. Five experimental conditions were used. These differed in the relative orientation of real and artificial hands (RH aligned to the body, or tilted by 90°), the relative hand position (besides or below the real hand), and one condition did not involve a rubber hand (Real condition). Each trial lasted 3 minutes and consisted of 180 visuo-tactile stimulation events of 100ms duration and presented with an interstimulus interval (ISI) was 900 ms, resulting in a visuo-tactile stimulation frequency of 1Hz. Visual stimuli were delivered by white light-emitting diodes (LED; Seeedstudio, 10 mm diameter) and tactile stimuli were delivered to the participant’s hand by a vibration motor (Grove: Vibration motor, Seeedstudio). Both stimuli were controlled via Matlab and two Arduino Uno prototyping platforms. To facilitate the alignment of stimulation events with the EEG data we routed a copy of the voltage controlling the DC motor to an analogue input of the EEG system.

The precise experimental conditions were designed as follows: In the ‘Illusion Hand Next’ condition, a lifelike rubber hand (for men: a silicon cosmetic glove, model 102LS, for women: model 102LS, ORTHO-REHA Neuhof GmbH) was positioned on top of the platform (horizontal plane) in front of the participants in an anatomically congruent orientation, as typically used to induce the illusion. The index finger of the rubber hand was placed on a dummy vibration motor, which did not vibrate. The participant’s left hand was covered with a blanket, hidden to the participant’s view, and was positioned at a distance of 10 cm from the rubber hand in the horizontal plane. The tip of the participant’s index finger was placed on the vibration motor, while the right hand was placed at the other end of the platform in reaching distance of a keyboard. The LED was positioned 5 mm above the dummy motor near the rubber hand. The somatosensory stimulus on the participant’s hand and the synchronous visual stimulus near the rubber hand reliably induce the illusion, as discussed below. In the ‘Control Hand Next’ condition, the rubber hand was placed in an anatomically incongruent position at a 90°angle. In the ‘Illusion Hand Under’ condition, the rubber hand was placed in front of the participant, while the participant’s real hand was covered with a blanket and placed in the lower panel of the platform 10 cm under the RH in the vertical plane. Otherwise the setup was the same as for the Illusion Hand Next condition. In the incongruent ‘Hand Under Condition’, the rubber hand was placed in an anatomically incongruent position, at a 90° angle below participants’ hand. We also included a control condition in which the rubber hand was absent. In this ‘Real’ condition no rubber hand was present, and the index finger of the left hand was placed on a vibration motor positioned 5 mm below the LED. The right hand was in the same position as in the Illusion and Incongruent conditions. In the following text we denote with the capitalized ‘Illusion’ the respective experimental conditions, while we use the non-capitalized illusion to refer to the phenomenon in general.

The choice of a rotated rubber hand for the control condition was made based on previous studies showing that an unrealistic posture abolishes the illusion reliably (Ehrsson, 2004; Pavani et al., 2000; Rao and Kayser, 2017; Riemer et al., 2019; Tsakiris and Haggard, 2005). Importantly, another often used control condition involving the asynchronous stimulation of the real and rubber hands can result in participants also reporting the illusion in presumed control conditions and makes specific assumptions about the temporal duration of the multisensory binding window, which the rotated hand avoids (Costantini et al., 2016; Fuchs et al., 2016; Valenzuela Moguillansky et al., 2013). Although some studies have shown that the mere observation of a rubber hand can at times also induce the illusion (Ferri et al., 2013; Tieri et al., 2015) our data suggests that this was not the case for the rotated control condition in the present sample of participants (see below). The Real condition was included as it allows comparing conditions with an embodied artificial and an embodied real hand, a dimension not probed by the misalignment of any real and artificial hands (Rao and Kayser, 2017; Zeller et al., 2016, 2015).

The choice of a computer-controlled stimulation setup using LED stimuli was made based on previous work. A previous study has shown that such a setup can reliably induce the RHI (Rao and Kayser, 2017). In particular, the LED and vibration stimulus provide a synchronized reference frame between the participant’s real hand and the rubber hand, which leads to the induction of the illusion (Bekrater-Bodmann et al., 2014; Evans and Blanke, 2013; Rao and Kayser, 2017; Shimada et al., 2009). Indeed, previous work has shown that the RHI can be reliably induced across different setups and manipulations that do not necessarily need to involve the presence of an experimenter stroking both hands with a brush (Bertamini and O’Sullivan, 2014; Ferri et al., 2013; Guterstam et al., 2015; Kalckert and Ehrsson, 2014). Furthermore, the brief and temporally precise stimulation provided by LED and the vibration motor allows the precise alignment of rapid neurophysiological signals to the stimulation sequence, which becomes intrinsically difficult with temporally imprecise events such as manual stroking (Rao and Kayser, 2017).

### Participants

To ensure that participants enrolled in the main EEG experiment were indeed able to feel the illusion, we conducted a pre-test. None of the participants tested in the pre-test reported having participated in a study involving the rubber hand or similar body illusions before. During the pre-test four conditions were presented in pseudo-random order (the two Illusion conditions and their Controls), each presented once. To probe whether or when participants felt the illusion we capitalized on question 3 from the common rubber-hand questionnaire (Table 1) (Botvinick and Cohen, 1998). Specifically, we instructed participants to press (using their right hand) a key on a computer keyboard when “feeling the rubber hand as belonging to their body”. After each trial, they were asked to confirm verbally that indeed they were feeling the rubber hand as belonging to their body. For the main study we invited only participants who in the pre-test had indicated feeling the illusion in both Illusion conditions and who did not report feeling the illusion in any Control condition. That only about half the pre-tested naive participants reported feeling the illusion fits with previous studies reporting a similar proportion (Ehrsson, 2004; Ehrsson et al., 2007; Lloyd, 2007; Riemer et al., 2019). All participants gave written informed consent before participation in accordance with the Declaration of Helsinki. All protocols conducted in this study were approved by the Ethics Committee of Bielefeld University.

**Table 1.** Questionnaire used to assess the subjective feeling of the illusion (adapted from Botvinick and Cohen, 1998). The questionnaire included nine statements describing the underlined phenomena. Participants indicated their response on a 7-scale ranging from “strongly agree” (−3) to “strongly disagree” (+3).

|  |  |  |
| --- | --- | --- |
| 1 | During the last trial it seemed as I were feeling the touch in the location where I saw the rubber hand touched | Illusion |
| 2 | During the last trial it seemed as though the touch I felt was caused by the vibration motor under the rubber hand | Illusion |
| 3 | During the last trial I felt as if the rubber hand were my hand | Illusion |
| 4 | During the last trial it felt as my (real) hand was drifting towards the rubber hand | Control |
| 5 | During the last trial it seemed as I might have more than one left hand or arm | Control |
| 6 | During the last trial it seemed as if the touch I was feeling came from somewhere between my own hand and the rubber hand | Control |
| 7 | During the last trial I felt as my (real) hand was turning rubbery | Control |
| 8 | During the trial it appeared (visually) as if the rubber hand were drifting towards left (towards my hand) | Control |
| 9 | During the trial the rubber hand began to resemble to my own hand in term of shape, skin tones, freckles or some other features | Control |

The main experiment took place on a different day than the pre-test. During the main experiment four repeats of each of the five conditions were administered in pseudo-random order for each participant. For each Illusion trial we determined the onset time at which participants started to feel the illusion as the time of the respective button press response. One participant was excluded from the main study as this participant reported feeling the illusion also during one Control trial and one participant was excluded due to reporting the illusion for only one Illusion condition. We also administered the full questionnaire (Table 1; Botvinick and Cohen, 1998) at the end of the first repeat of each of the two Illusion conditions. From these we calculated the average score for illusion (Illusion hand next: 2.38 ± 0.53, mean ± s.e.m.; Illusion hand under: 2.61±0.53) and control statements (hand next: - 1.47 ± 0.76; hand under: -1.56 ± 0.80).

### EEG recording and pre-processing

EEG signals were continuously recorded using a 128 channel BioSemi (BioSemi, B.V., Netherlands) system with Ag-AgCl electrodes mounted on an elastic cap (BioSemi). Four additional electrodes were placed at the outer canthi and below the eyes to obtain the electro-oculogram (EOG). Electrode offsets were kept below 25 mV. Data were acquired at a sampling rate of 1028 Hz. Data analysis was performed with MATLAB (The MathWorks Inc., Natick, MA, USA) using the FieldTrip toolbox (Oostenveld et al., 2010). Data were band-pass filtered between 0.4 Hz and 90 Hz, resampled to 200 Hz. Subsequently, the data were denoised with ICA. Components reflecting muscular artefacts, eye blinks, eye movements as well as poor electrode contacts were identified based on recommendations in the literature (Hipp and Siegel, 2013; O’Beirne and Patuzzi, 1999) were removed similar to our previous work (Bröhl and Kayser, 2021; Grabot and Kayser, 2020). Overall, we removed an average of 17.45 ± 1.48 (mean ± s.e.m.) components per participant.

For subsequent data analysis we epoched the data around each visuo-tactile stimulation event, with epochs lasting from 250 ms pre-stimulus to 400 ms post-stimulus onset This resulted in a total number of 180 epochs for each trial. Epochs were combined within each condition, were band-pass filtered between 2 Hz and 40 Hz, and epochs with signals exceeding a level of 250 µV were removed. Note that we used such a lenient threshold as the classification analysis employed here is rather insensitive to outliers. Importantly, to render the analysis of the Illusion conditions specific to participants experiencing the illusory state, we only included epochs after the time point at which participants indicated the onset of the illusion. Hence for any analysis of the Illusion conditions, we included only date epochs after the respective onset of the subjective illusion state. As a result, the number of epochs available for each condition differed. The average number of epochs for both Illusion conditions was 1347 ± 34 (mean ± s.e.m.), for the Control conditions 1299 ± 55, and for the Real condition 684 ± 15. To avoid biasing analyses by these differences in epoch numbers we used sub-sampling of epochs to equalize the number of epochs per condition for each statistical contrast described below. Sub-sampling was based on randomly selecting (without replacement) 80% of the minimally available number of epochs for each contrasted condition, and repeating each calculation multiple times using random selections of epochs.

### Analysis of EEG activity using single-trial classification

To quantify whether and when EEG activity differed between experimental conditions, we used single-trial classification based on a regularized linear discriminant analysis (LDA) (Blankertz et al., 2011; Parra et al., 2005). The epoched data were binned into overlapping time bins of 60ms duration and the classifier was computed at 10 ms time steps. Results are reported aligned to the onset of the 60ms window. The regularization parameter was set to 0.1 as in previous work (Park and Kayser, 2019). For each contrast we combined the epochs from the respective trials of interest, using the sub-sampling procedure described above (e.g. all Illusion vs. all Control epochs; or only the hand next illusion vs. only the hand next control epochs). For each participant the classifier performance was obtained as the receiver operating characteristic area under curve (AUC), computed from 6-fold cross-validation. That is, we trained the data on five-sixth of the data and tested the classification performance on the remaining one-sixth of the data. We derived participant-wise scalp topographies for each classifier by estimating the corresponding forward model, defined as the normalized correlation between the discriminant component and the EEG activity (Parra et al. 2005). Importantly, by combining the signals from all electrodes for the classifier this analysis does not make the assumption that the condition-wise spatial configuration of activity is the same across participants. This classification analysis was used to test for EEG responses differentiating i) the Illusion vs. Control epochs (across both hand positions; and separately for each hand position), ii) the Illusion vs. Real epochs, iii) the effect of hand position regardless of illusory state (by grouping Illusion and Control epochs according to hand next and hand under labels), iv) and for activity differentiating those epochs within the Illusion trial prior to the onset of the illusory state (hence before participant’s button press) versus those during the illusion (i.e. after the button press).

To directly test whether the neurophysiological signatures discriminating e.g. the Illusion and Control conditions are the same as those also differentiating the Illusion and Real conditions, we used cross-classification. For this we trained the classifier on half the Illusion and the Control trials and then applied classifier weights to differentiate the remaining half of the Illusion and the Real trials, repeating this procedure 50 times. Cross-classification was again quantified using the AUC and computed for both directions: training on Illusion vs. Control and testing on Illusion vs. Real, and training on Illusion vs. Real and testing on Illusion vs. Control. The respective AUC values for each participant and time point were averaged over both directions and repeats of the calculation. The same cross-classification analysis was used in the following further analyses: i) to test whether the neurophysiological signatures discriminating Illusion and Control for the ‘hand next’ arrangement generalize to the ‘hand under’ arrangement, and vice versa, ii) to test whether activity differentiating the overall hand position (all ‘hand next’ vs. all ‘hand under’ epochs) also differentiates Illusion from Control epochs, iii) and to test whether classifiers trained on the discrimination of Illusion vs. Control (or Illusion vs. Real) also generalize to the discrimination of the epochs prior to the Illusion vs. those during the Illusion.

### Recording of skin conductance signals

Skin conductance was continuously recorded using a Neulog GSR logger NUL-217 sensor with Ag/AgCl electrodes were placed on the palmar sites of the middle and ring finger of the participant’s left hand. GSR measurements were recorded in micro-Siemens and at a sampling rate of 100 Hz. For offline analysis the data were resampled to 50 Hz, band-pass filtered between 0.5 Hz and 2 Hz (3rd-order Butterworth filter) and logarithmically (Boucsein et al., 2012; Sjouwerman and Lonsdorf, 2019) (Braithwaite et al., 2013). Trials were visually inspected and few trials with artefacts were manually removed. To determine whether the changes in skin conductance observed in the main experiment could be the result of simply pressing a button (vs. not pressing any button) we also recorded skin conductance data in a separate control session in which participants only had to press a button in response to a visual cue for eight times. For a few participants (n=5) the skin conductance data were incomplete, as the software had crashed during recording. Hence the effective participant sample for EEG and skin responses differed.

### Analysis of skin conductance data

We opted for an analysis that directly contrasts the skin conductance in two adjacent time windows of 3 s duration (Boucsein et al., 2012). These windows were chosen around the key event of interest: the trial-specific onset of the illusion indicated by participants pressing the response button. The duration of the analysis window was chosen as a compromise between the typical latency of changes in skin conductance (Boucsein et al., 2012; Sjouwerman and Lonsdorf, 2019) and the reaction times at which participants started feeling the illusions (c.f. Fig. 2). To quantify the change in skin conductance (termed ‘skin response’) induced by the illusion onset we computed the standard deviation (SD) of the signal in two adjacent windows around the illusion onset (e.g. [-3 s, 0 s] and [-0 s, +3 s]) and then computed their ratio. In one analysis we computed the ratio between the window immediately before the illusion onset ([-3 s, 0 s]) and the immediate following window ([0 s, +3 s]). In a second analysis we focused on the ratio of the two windows immediately prior to the illusion onset (hence [-6 s, -3 s] and [-3 s, 0 s]). To calculate the standard deviation (and then, their ratio) for Control and Real conditions, we used trial-number-specific (i.e. 1-4) reaction times obtained from the Illusion trial with that respective number as surrogate time points to define the corresponding windows. For the button press session, we extracted the skin conductance in time windows aligned on the button press time.

### Statistics

Statistical testing of the classification performance (AUC values) relied on a randomization approach and cluster-based permutation procedures to exploit the smoothness of the data and to control for multiple comparisons (e.g. across time) (Maris and Oostenveld, 2007; Nichols and Holmes, 2002). In general, we relied on 2000 randomizations for each test, used a cluster-forming threshold corresponding p<0.01 (i.e. using the 99th percentile of the full distribution of randomized AUC values), applied spatial clustering based on a minimal cluster size of 3, and used the sum of AUC values within each cluster as cluster-wise test statistics. The randomization distribution was obtained by shuffling the sign of the true single-participant effects (i.e. the sign of the chance-level corrected AUC values) based on which we derived a distribution of expected group-level effect of no systematic classification performance across participants. For significant clusters we report the cluster statistics (summed AUC value) and the peak classification performance (max AUC). To compare the median latencies of illusion onsets between conditions and to compare the skin responses between pairs of conditions we used Wilcoxon signed rank tests. Effect sizes for this test were obtained using the point-biserial correlation (Kerby, 2014).

## Results

### Illusion onset times

We compared the mean onset times of the illusory states between the two Illusion conditions. The onsets in the ‘hand next’ condition occurred on average after 50 ± 35 s (n=22; mean ± SD; median: 42 s) and significantly later compared to the ‘hand under’ condition (34±27 s, mean ± SD; median: 32 ms; Wilcoxon signed rank test Zval=1,96, p=0.04, r_rb_ = 0.41). Hence, the hand under arrangement required less time to induce the illusion, despite the physical distance between the participant’s and the rubber hand being the same in both configurations.

### Skin conductance

We implemented two analyses geared to reveal changes in skin conductance emerging either in parallel with the participant’s overt response of feeling the illusion (button press), or prior to this. To remove slow drifts in the recorded signals, the analysis was based on the ratio of the skin conductance in two subsequent 3s windows (termed ‘skin response’), either centered on the button press or an epoch preceding this. The analysis centered on the response times revealed significant differences in skin response between Illusion and Control conditions (n= 17, Wilcoxon sign-rank test, Z=2,9, p_corr_ = 0.02, r_rb_=0.80; Figure 1B) but not between Illusion and Real conditions (n=17, Z=1,59, p_corr_ =0.22, r_rb_=0.43). We also did not find a significant difference between the Illusion condition and a separate control experiment in which participants simply pressed a response button (p_corr_ =0.22, r_rb_ = 0.46). The analysis focusing on the epochs prior to the overt response revealed significantly stronger skin responses during the Illusion compared to the Control condition (n=17, Z=3.29 p_corr_ =0.003, r_rb_=0.90), but again not between the Illusion and Real condition (n=17, Z=1.54, p_corr_ =0.18, r_rb_=0.42).

**Figure 1.**
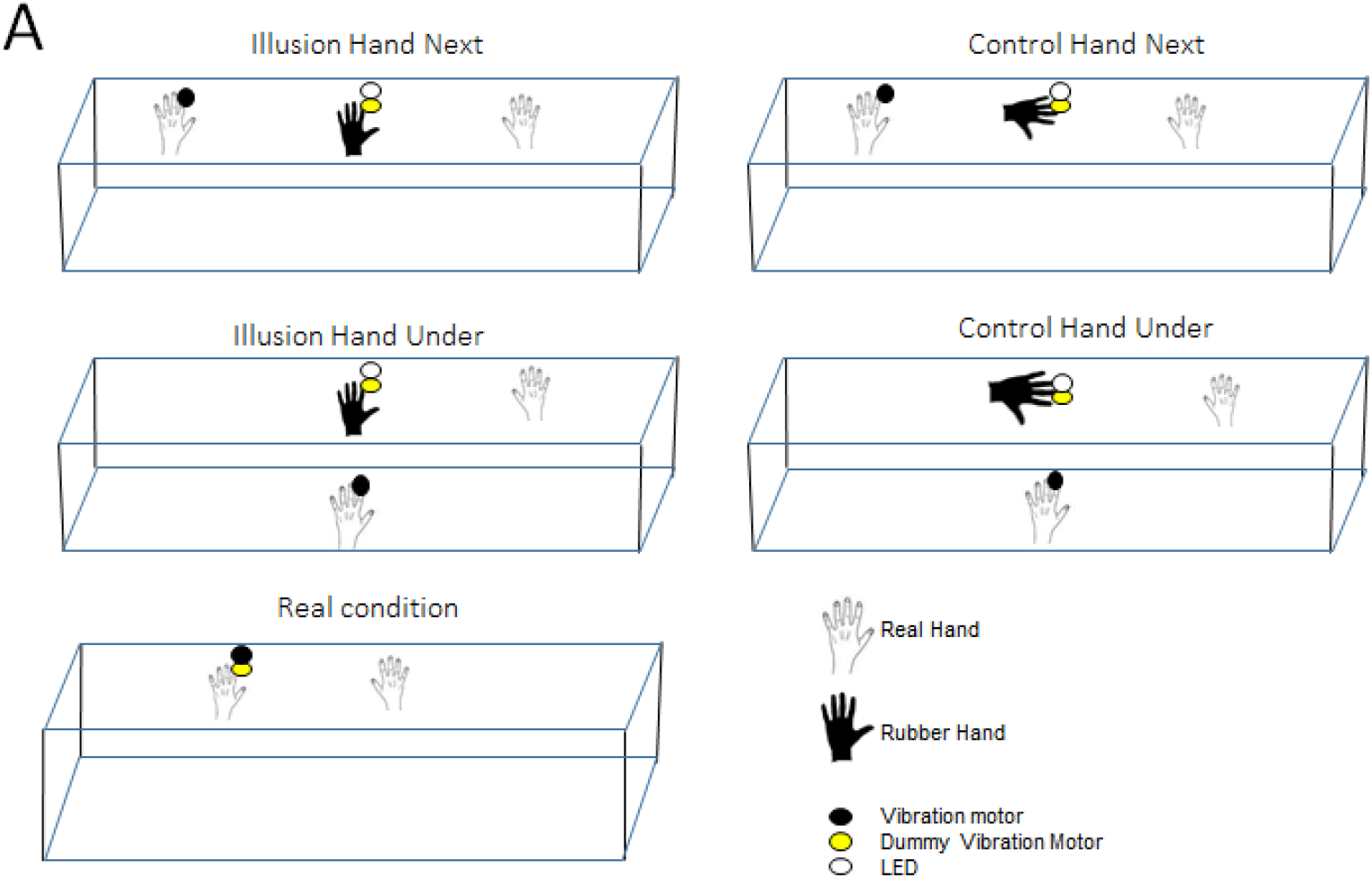
Schematic of the five experimental conditions. The experiment involved two ‘Illusion conditions’ that differed in the spatial arrangement of the real and artificial limbs (horizontal or vertical displacements), and the two respective ‘Control conditions’ obtained by rotating the rubber hand into an unphysiological position by 90 degrees. Finally, we included a ‘Real condition’ in which the rubber hand was absent and multisensory stimulation was delivered to the real hand.

### Evoked responses differ between Illusion and Control conditions at short latencies

We then asked whether and how brain activity differs between epochs in which participants feel the illusion (i.e. the illusory-state) and the different non-illusion conditions. Specifically, we focused on the electrophysiological response evoked by the repetitive visuo-tactile stimulus as a signature of the cerebral processing of the relevant distant signals. To quantify potential changes between conditions, we relied on a multivariate linear discriminant analysis. This avoids the assumption implicitly made in previous studies (Cichy and Pantazis, 2017) Aspell et al., 2012, Peled et al., 2003, Rao and Kayser, 2017; Zeller et al., 2015) that the illusion-related somatosensory evoked processes have the very same spatial topography across participants. Using a multivariate classifier avoids this assumption by allowing distinct spatial patterns of activity to contribute to condition-differences in individual participants, while assuming that any systematic group-level differences between conditions emerge around the same time in the data.

Group-level classification performance of Illusion versus Control conditions was significant from 0.01 s to 0.320 s after stimulus onset (n=22, Cluster-based permutation test correcting for multiple comparisons along time, p<0.001, AUC_sum = 3.68, max AUC = 0.69 at 0.100 s; Fig. 2A) suggesting that evoked activity differs between conditions at early latencies following the illusion-inducing stimulus. In a subsequent analysis we confirmed that the difference between Illusion and Control conditions prevailed for both relative positions of the artificial and real hands: classification of each Illusion versus the respective Control condition was significant in similar time windows (hand next: from 0.01 to 0.320 s, p<0.001, AUC_sum=3.67, max AUC=0.73 at 0.100 s; hand under: from -0.01 s to 0.300 s, p<0.001, AUC_sum=3.20, max AUC=0.68 at 0.100 s; Fig. 2B). Contrasting the Illusion and Real conditions revealed a similar widespread difference (n=22, cluster from 0.04 s to 0.310 s, p<0.001, AUC_sum=2.64, max AUC=0.66 at 0.90 s; Fig. 2A), suggesting that the illusory state affects evoked response compared to both types of non-Illusion conditions.

**FIGURE 2.**
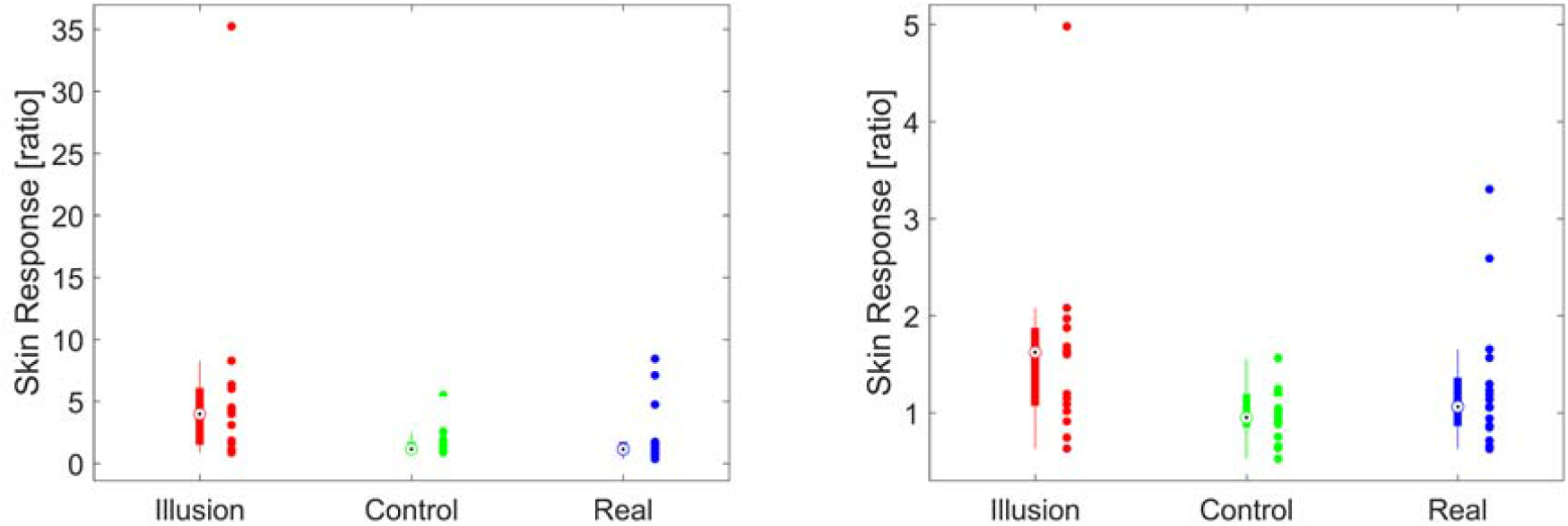
Changes in skin conductance associated with the illusion. Boxplots show skin responses around the reported onset of illusion (upper panel) and in a 3s period prior to the reported onset of the illusion (lower panel). Skin responses were defined as the ratio of the conductance in a 3s window of interest, normalized to a preceding 3s baseline. Boxplots indicate the median (circle), 25th and 75th percentiles (thick line). Dots indicate individual participants.

We then asked whether the differences between the Illusion and Control conditions and between Illusion and Real conditions arise from the same neurophysiological processes and hence have a similar spatial configuration. To address this question we relied on the concept of cross-decoding (Cichy and Pantazis, 2017). Using cross-decoding one can establish a signature of similarity between experimental conditions, here in the form of a spatio-temporal configuration of electrode-wise activity contributions, which are reflected by the electrode-specific classifier weights. Based on these, one can ask whether these configurations generalize across conditions. Practically, this is tested by training the classifier to discriminate between one pair of conditions (e.g. Illusion vs. Control) and testing this on another pair (e.g. Illusion vs. Real), using non-overlapping sets of trials.

The cross-decoding analysis was significant between 0.130 s to 0.150 s (p<0.001, AUC_sum=0.08, max = 0.53 at 0.130 s; Fig. 2C), suggesting that during these time points the illusion epochs are distinguished from both types of non-illusion epochs by a similar pattern of evoked response. We then used the same cross-decoding approach to investigate whether the neurophysiological processes differentiating the Illusion and the Control for the ‘hand next’ conditions shared a similar spatial configuration compared to processes differentiating these conditions during the ‘hand under’ configuration. Cross-decoding was significant for most time points (cluster 1: from 0.05 s to 0.120 s, p<0.001, AUC_sum=0.35; cluster 2: from 0.150 s to 0.190 s, p<0.001, AUC_sum=0.17; cluster 3: from 0.250 s to 0.270 s, p<0.001, AUC_sum=0.09; max AUC= 0.55 at 0.09 s; Fig. 2D). Together, these cross-decoding results suggest that the neurophysiological processes differentiating the Illusion from the two Control conditions fall into two types: one that generalizes across non-illusion conditions (Control and Real) from 130 ms to 150 ms following the visuo-tactile stimulus, and others for which activity selectively differentiates the Illusion from either the control conditions involving the rotated limb (Control) or a condition only involving participants real hand (Real).

### Evoked responses differ within trials according to the subjective illusion

Statistical comparisons between Illusion and Control conditions are usually performed by comparing all epochs from the Illusion condition against all those from the Control conditions. However, the experimental Illusion and Control conditions are provided at different times during the experiment and hence may differ not only in the subjective illusory state. To make the comparison of illusion and non-illusion related brain activity more specific, we directly contrasted epochs characterized by the illusory-state and those reflecting non-illusion epochs within the same experimental Illusion trials: based on participants’ responses indicating the onset of the illusory state, we split the epochs from the Illusion trials into those prior to the illusion onset and those during illusory state: the classification analysis between these was also statistically significant (n=22, cluster from 0.03 to 0.300s, p<0.01, AUC_sum=2.11, max AUC=0.65 at 0.130s; Fig. 3A).

**Figure 3.**
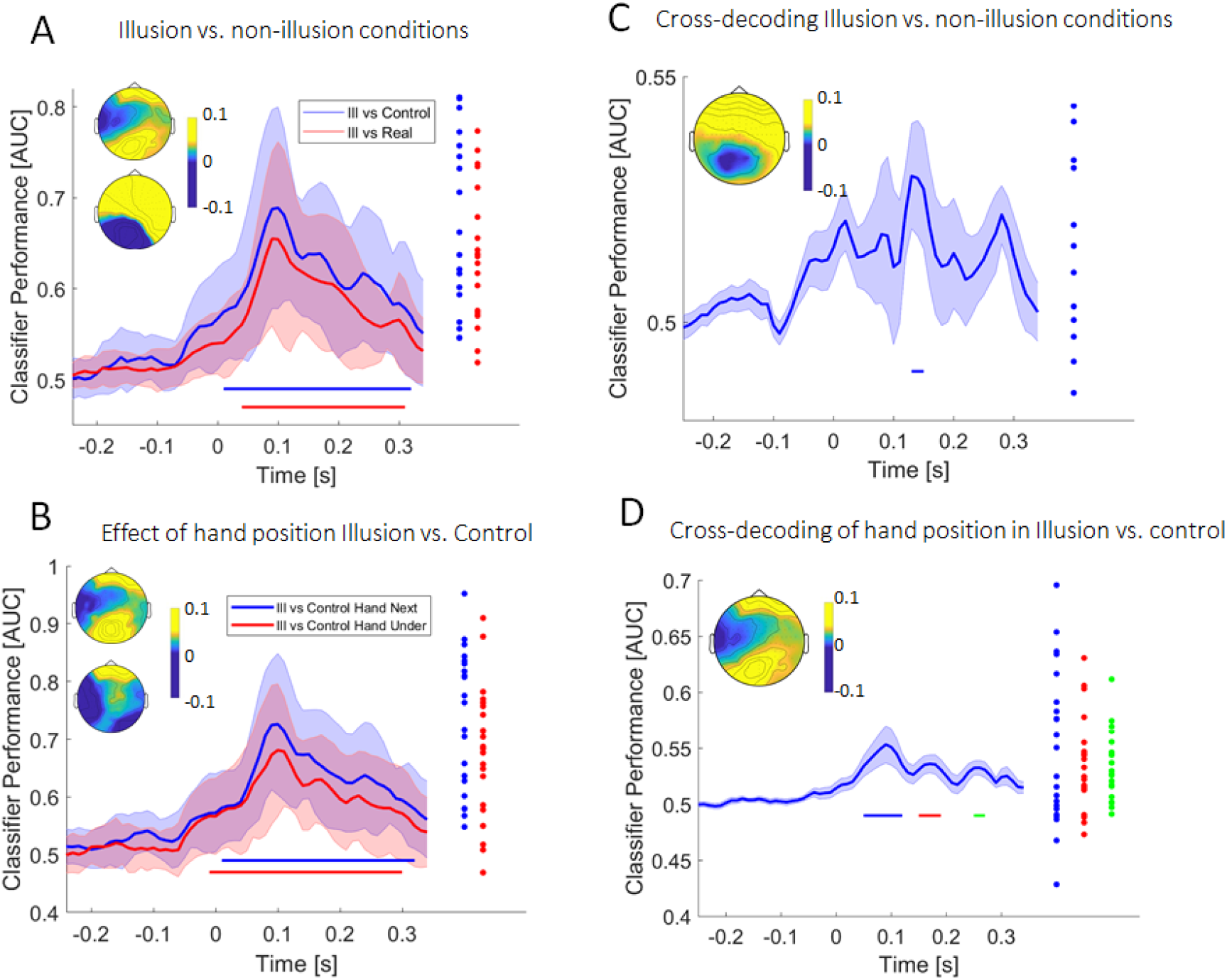
Differences in EEG evoked responses associated with the illusion. **A)** Differences in EEG activity between Illusion and Control conditions and between Illusion and the Real condition quantified using linear multivariate classification. Classifier performance was computed as the area under the receiver-operator characteristic (AUC). Thick curves indicate the mean and shaded areas indicate the s.e.m. Straight lines at the bottom indicate periods of significant classification performance (cluster-based permutation test corrected for multiple comparisons along time, at p<0.01). Topographies indicated the spatial distribution of the activity at the time points of maximal classifier performance. Group-level forward models of the classifier weights at the time of the maximal classification performance. Upper right: Illusion vs Control. Lower right: Illusion vs Real. Dots indicate individual participants classifier performance. **B)** Same results for a classification between each individual Illusion condition and the respective Control condition. Group-level forward models of the classifier weights at the time of the maximal classification performance. Upper right: Illusion vs Control for hand next conditions. Lower right: Illusion vs Control for hand under conditions. **C)** Performance of a cross-decoding analysis between Illusion vs. Control and Illusion vs. Real conditions. The classifier was trained on one of these contrasts and tested on the other, with the result averaged over both directions. **D)** Performance of a cross-decoding analysis between each individual Illusion and its respective Control condition.

**Figure 4.**
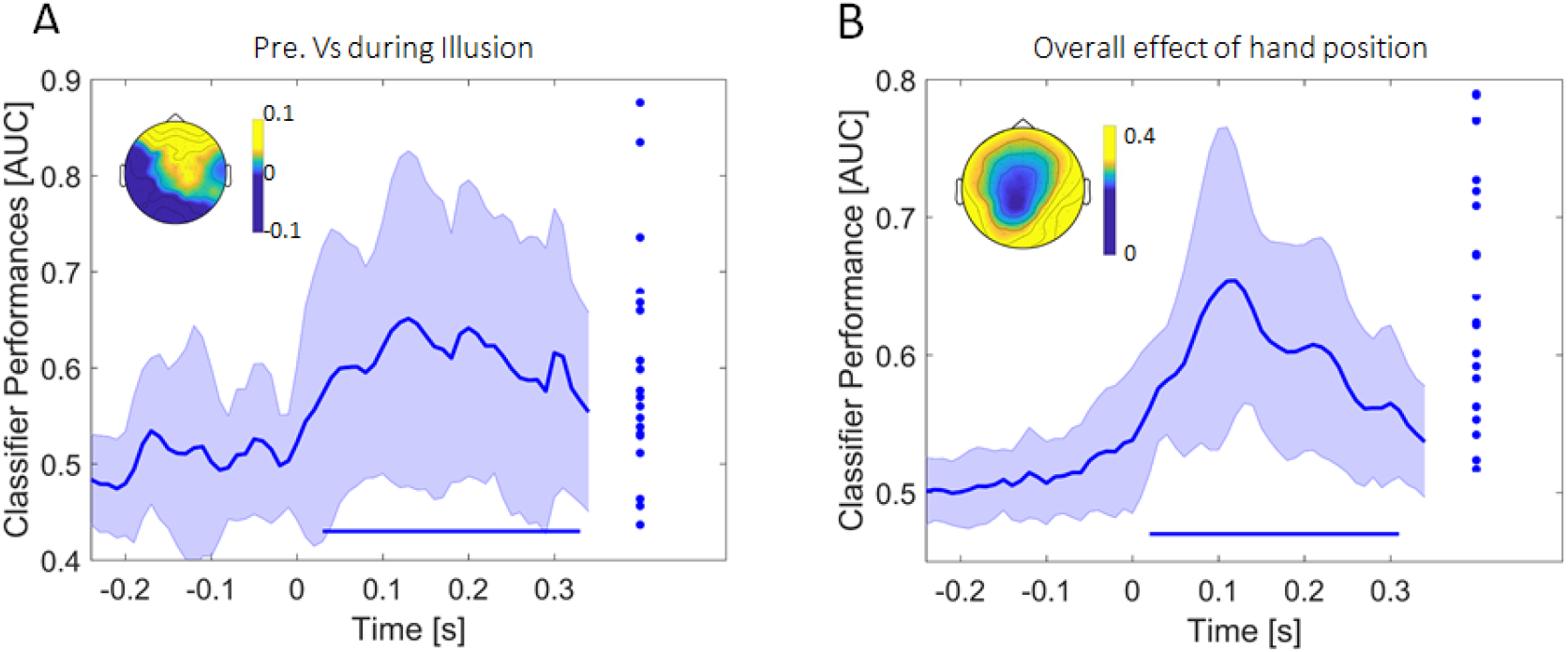
Differences in pre-illusion vs. during-illusion epochs within a trial, and effect of hand position. **A)** Classification of EEG activity between the epochs before the illusion onset and the epochs after the illusion onset within the Illusion trials. Classifier performance is shown as the area under the receiver-operator characteristic (AUC). Thick curves indicate the mean and shaded areas indicate the standard errors of the mean. Straight lines at the bottom indicate periods of significant classification performance (cluster-based permutation test correcting for multiple comparisons along time, at p<0.01). Scalp topographies show the classifier forward models at the time of the maximal classification performance. Dots are the individual Classifier performance at the time point of maximal performance) **B)** Main effect of hand position. Classification performance (AUC) for ‘hand next’ versus ‘hand under’ epochs regardless of illusory state. Thick curves indicate the mean and shaded areas indicate the standard errors of the mean. Straight lines at the bottom indicate periods of significant classification performance (cluster-based permutation test correcting for multiple comparisons along time, at p<0.01).

We then asked whether the activity patterns differentiating the pre-illusion and Illusion epochs within the same experimental trials are similar to those differentiating the Illusion and Control epochs (Control, Real). To address this, we trained a classifier on the pre-illusion vs. Illusion classification and tested whether this generalizes to the discrimination of Illusion vs. Control (or Illusion vs. Real) epochs. Neither of these cross-classification analyses was significant (cluster-based permutation tests, no clusters at the respective criteria; max AUC=0.52 at 0.300s).

### Effect of hand position

The use of two spatial arrangements to induce the illusion allowed us to separate the effect of the illusory state from a potential influence arising from the relative positioning of real and artificial hands. While our results show that the two Illusion conditions differed in their efficacy in inducing the illusory state (the ‘hand under’ condition led to a more rapid onset of the illusion) the cross-classification analysis confirmed neurophysiological processes that reflect the illusory state regardless of the spatial configuration. In an additional analysis we then asked whether brain activity also differs with the main effect of hand position, hence between the two conditions (Illusion and Control) featuring the participants real hand *next* to the rubber hand and the two conditions featuring the participants hand *under* the rubber hand. While we expect such processes to exist, the above results suggest that these should be independent of those characterizing the illusory state.

We tested these questions using the following classification and cross-classification analyses. First, we probed classification performance between conditions featuring the ‘hand next’ and ‘hand under’ arrangements: this was significant from 0.02 s to 0.310 s (n=22, p<0.01, AUC_sum=3.02, max AUC=0.65 at 0.120 s; Fig. 3B). Second, we used cross-decoding to test whether the spatial configuration of the EEG activity discriminating hand positions also discriminates the effect of illusion, defined by the contrast Illusion vs. Control conditions. Cross-classification between the effects of illusion and hand position was not significant (cluster-based permutation test, no clusters at the respective criteria, max AUC = 0.51).

### Neurophysiological illusion signatures are not related to skin responses

Given that the illusory state was associated with changes in EEG evoked responses and with changes in skin conductance, we asked whether the respective effects were correlated across participants. To test this, we computed the correlation between the LDA classifier performance in discriminating Illusion and Controls trials as a measure of illusion-effect in the EEG signal and the skin responses in the Illusion conditions as a measure of autonomic response. To assess the significance of these correlations we derived group-level bootstrap confidence intervals. For neither time window for skin response and neither LDA classifier was the correlation significant at any time point (all p>0.05 two-sided; skin response at illusion onset: LDA Illusion vs. Control had maximal r^2^ = 0.25 at any post-stimulus time of 0.16; LDA Illusion vs. Real r^2^ = 0.20; LDA After Vs Before r^2^=0.03; skin response prior to button press: r^2^=0.004, 0.34 and 0.23 respectively).

## Discussion

The aim of this study was to investigate the physiological correlates of the illusory state during the RHI. We used multivariate (cross-)classification of evoked EEG responses to ask whether and when these differentiate the illusory state reliably from several non-illusory control conditions involving different relative positions of both hands or the stimulation of the participant’s real hand. We found that evoked responses differed between illusion and non-illusion epochs already at short latencies following stimulation onset and over prolonged time. These neurophysiological signatures were not related to changes in skin conductance that accompanied the onset of the illusory state.

### Correlates of the illusory state in evoked responses

A number of studies aimed to understand the EEG-derived correlates of the RHI by investigating the responses evoked by the illusion-inducing stimuli (Aspell et al., 2012, Cardini et al., 2011, Peled et al., 2003; Press et al., 2008, Rao and Kayser, 2017, Zeller et al., 2015). This approach rests on the idea that the neurophysiological signatures in the evoked response can reliably index the relevant neurophysiological processes differentiating the illusory state from suitable control conditions. However, these previous studies described conflicting results: while some reported a modulation of evoked responses at early (55ms) latencies by the illusion (Otsuru et al., 2014; Zeller et al., 2015) others reported effects at latencies around 140-200 ms (Press et al., 2008) or even later, such as between 350 and 450 ms (Peled et al., 2003, Rao and Kayser, 2017). Importantly, all studies investigating group-level effects in evoked responses made the implicit assumption that the neurophysiological sources differentiating illusion and non-illusion conditions have the same spatial configuration across participants. We exploited a multivariate classification framework that allowed us to relax this assumption. Importantly, the classification analysis allowed us to directly test, within each participant, at which time point the relevant neurophysiological signatures generalize between conditions, such as the Control and Real conditions, or horizontal and vertical arrangements of real and artificial hands. Our results show that evoked responses differ reliably between illusion and non-illusion epochs over prolonged periods and at short latencies.

The earliest differences between illusion and non-illusion epochs emerged shortly after stimulus onset, and hence likely arise from early somatosensory cortices. These condition differences did not generalize between different non-illusion conditions or between hand positions, in line with the underlying primary sensory regions encoding information in a highly specific manner. Subsequently, between 50-100 ms, the illusion-related processes generalized across hand positions in the Illusion vs. Control contrast, suggesting that the respective processes are invariant to relative body arrangements. However, the activity patterns did not generalize to the Real condition, possibly because the underlying cerebral processes arise from somatosensory cortices that are sensitive to the bodily nature of the seen hand (Aspell et al., 2012, Zeller et al., 2015, Cardini et al., 2011, Cardini et al, 2012, Otsuru et al, 2014). Such early differences in evoked EEG activity between the illusory state and control conditions have been reported before, but these studies did not directly test whether and which illusion-correlates generalize between factors.

Importantly, in our study the difference between illusion and non-illusion epochs generalized across all control conditions between 130 and 150 ms, making activity around this time a robust marker for the illusory state. Supporting this, Press et al., 2008 found an enhanced somatosensory N140 after synchronous versus asynchronous stimulation, irrespective of whether the participants viewed a rubber hand or object. Together with the present data, this suggests that systematic and consistent differences between illusion and multiple control conditions arise around these latencies. The forward models of the classifiers (c.f. Fig. 2) revealed a differential contribution of bilateral fronto-parietal sensors, as opposed to for example a lateralization of the relevant processes. Hence, this neurophysiological evidence is consistent with the notion that parietal or premotor regions integrate the visual-tactile information about body position and underpin the sensation of ownership (Blanke, 2012; Ehrsson, 2004; Fang et al., 2019; Graziano et al., 1994, 2000). Still, the present data cannot unequivocally determine the precise origin of the underlying neurophysiological processes. Future studies, for example relying on combined EEG-fMRI recordings may capitalize on the present experimental and analytical approach to more precisely determine the brain regions correlating with, or even giving rise to the illusion. In particular, such experiments could provide insights into what aspects of the illusory setting (e.g. spatial attention, seeing an artificial body, feeling ownership over an object) affect the neurophysiological representation of the sensory stimulation at distinct stages of somatosensory or interoceptive pathways. Also, we did not administer another typical control condition provided by the asynchronous stimulation of participant’s and the rubber hand, and it remains to be seen whether these evoked responses are sensitive or not to such temporal incongruency.

Lastly, after about 160ms, differences between the illusion and non-illusion conditions generalized across hand positions, but remained specific to the Control or Real condition. One possibility is that these effects reflect higher parietal and frontal processes sensitivity to multiple factors, such as spatial attention (Rao and Kayser, 2017), bodily self-detection (Galigani et al., 2021) or high-level processes pertaining to multisensory causal inference (Cao et al., 2019; Rohe and Noppeney, 2015). More work is required to understand the precise neurophysiological origins and what precise aspect of the experimental paradigm or subjective state these are selective to.

A parallel body of neurophysiological studies has focused on rhythmic brain activity and its relation to body illusions. These studies reported putative illusion-correlates in alpha, beta or gamma band oscillations or between-electrode synchronization (Faivre et al., 2017; Kanayama et al., 2021, 2009, 2007; Lenggenhager et al., 2011; Shibuya et al., 2019). However, given the inherent difficulties to precisely localize oscillatory activity in time, these indices of rhythmic activity are less well suited to address the question of when following the visuo-tactile stimulation events illusion-specific correlates emerge, the main question addressed here.

### The impact of methodological details on outcomes in studies on the RHI

In studies on the RHI the Illusion and Control conditions generally differ along multiple dimensions. These include aspects such as the subjective state of body ownership, the relative positions or configurations of body parts, the temporal pattern of sensory stimuli, or the nature of the presented control objects (Bekrater-Bodmann et al., 2014; Bertamini and Sullivan, 2014; Ferri et al., 2013; Guterstam et al., 2013; Kalckert and Ehrsson, 2014; Rao and Kayser, 2017; Shimada et al., 2009; Tsakiris and Haggard, 2005). This makes it difficult to associate neurophysiological correlates of the illusory state with one specific aspect of this paradigm. In a step to overcome this problem, we here combined multiple control conditions with multivariate cross-decoding, which allowed us to directly probe which neurophysiological processes consistently differentiate the illusory state from more than one control condition.

First, we employed two spatial configurations of the real and rubber hands for the Illusion conditions, by displacing the RH and real hands either in the horizontal or vertical planes, while keeping their physical distance the same. Our results showed that the hand-under arrangement required less time to induce the illusion, in line with studies reporting stronger illusory precepts for the vertical set-up based on questionnaire scores (Bekrater-Bodmann et al., 2012) and with the general idea that distances in the horizontal and vertical plane are often judged differently (Loomis and Philbeck, 1999). Second, we employed two distinct experimental control conditions as contrasts for the illusion and a further, within trial, control analysis. The synchronous stimulation of a bodily-misaligned rubber hand, as used in the Control conditions, has proven more effective than the temporally asynchronous stimulation of a body-aligned rubber hand (Costantini et al., 2016; Fuchs et al., 2016; Riemer, 2019; Valenzuela Moguillansky et al., 2013). In addition, we used participants’ real hand as a control, a condition in which the rubber hand was absent and the visuo-tactile stimulation directly occurred on the participant’s (naturally embodied) real hand. And finally, we directly contrasted the activity within the Illusion trials between periods before and during participants feeling the illusion. This contrast pertains only to the Illusion configurations and allows a comparison with the very same sensory stimulation and configuration, with the only difference supposedly being the participant’s subjective illusion.

The cross-decoding approach revealed neurophysiological signals differentiating illusory and non-illusory epochs (Control, Real) that generalize across the spatial plane in which the illusion is induced and between control conditions involving a rotated rubber hand or participants’ own hand. However, those neurophysiological processes did not allow differentiating the epochs prior to and following the onset of the illusory state in the Illusion trials. This provides evidence that distinct neurophysiological processes differentiate between distinct experimental conditions and the subjective state within the same experimental condition. While we can’t rule out that the pre vs. during illusion contrast was affected by the small number of pre-illusion data epochs available for some of the participants, it will be interesting to better characterize the neurophysiological changes emerging directly at the time participants begin to feel the illusion, given that this contrast can rule out a number of confounding factors present when contrasting distinct experimental configurations. However, this within-trial comparison is also susceptible to adaptation effects due to the repetitive stimulation of the somatosensory system (McLaughlin and Kelly, 1993), and such adaptation effects may potentially confound with presumed illusion correlates in this specific analysis.

### Changes in skin conductance associated with the onset of the illusion

Previous studies have reported that the RHI is associated with changes in bodily arousal indexed by skin conductance (Armel and Ramachandran, 2003; Ehrsson et al., 2007). Often, a threat is applied to the embodied RH, which induces changes in skin conductance compared to control conditions (Armel and Ramachandran, 2003; Ehrsson et al., 2007, Petkova and Ehrsson, 2012). However, the threat may induce hand movements which can bias skin conductance results (Kilteni et al., 2012) and increased skin conductance after a threat has also been found independently of embodiment (Riemer et al., 2015). To overcome those problems, recent work began to investigate changes in skin conductance during the entire experimental trial (D’Alonzo et al., 2020).

Following this approach, we focused on the moments at which the illusory state emerged and asked whether this emergence is characterized by concomitant changes in skin conductance. Our data support a change in bodily state prior and around participants’ actual overt response of reporting the illusion, possibly because changes in arousal accompany the subjective sensation of ownership. The associated increase in skin response was significant in comparison to the Control but not when compared to the Real condition. We take this to suggest that these autonomous responses are most strongly modulated by seeing a not-embodied object rather than by the nature of the embodied object, be it a real rubber or a real hand.

While bodily signals such as skin conductance are frequently investigated in relation to the RHI and their suitability as markers of the illusory state remain debated (Crucianelli et al., 2018; Horváth et al., 2020; Suzuki et al., 2013). In particular, whether and to what degree bodily and neurophysiological markers of the illusion reflect the same underlying processes remains unclear, as some studies also reported dissociations between various signatures of the illusion such as proprioceptive drift and questionnaires (Holle et al., 2011; Holmes et al., 2012; Horváth et al., 2020; Kammers et al., 2011; Riemer et al., 2015). We did not find a significant correlation between skin responses and the degree to which the evoked activity differed between illusion and non-Illusion conditions. Hence, our data rather support the notion that both types of changes in physiological state are driven independently and characterize distinct processes contributing to the subjective feeling of the illusion.

## Acknowledgements

CK was supported by the European Research Council (ERC-2014-CoG; grant No 646657). We are grateful to Alexander Wecker for help with implementing the experimental setup.

## Bibliography

Armel, K.C., Ramachandran, V.S., 2003. Projecting sensations to external objects: evidence from skin conductance response. Proceedings of the Royal Society of London. Series B: Biological Sciences 270, 1499–1506. https://doi.org/10.1098/rspb.2003.2364

Aspell, J.E., Palluel, E., Blanke, O., 2012. Early and late activity in somatosensory cortex reflects changes in bodily self-consciousness: An evoked potential study. Neuroscience 216, 110–122. https://doi.org/10.1016/j.neuroscience.2012.04.039

Bekrater-Bodmann, R., Foell, J., Diers, M., Kamping, S., Rance, M., Kirsch, P., Trojan, J., Fuchs, X., Bach, F., Çakmak, H.K., Maaß, H., Flor, H., 2014. The Importance of Synchrony and Temporal Order of Visual and Tactile Input for Illusory Limb Ownership Experiences – An fMRI Study Applying Virtual Reality. PLoS ONE 9, e87013. https://doi.org/10.1371/journal.pone.0087013

Bertamini, M., O’Sullivan, N., 2014. The use of realistic and mechanical hands in the rubber hand illusion, and the relationship to hemispheric differences. Consciousness and Cognition 27, 89–99. https://doi.org/10.1016/j.concog.2014.04.010

Blanke, O., 2012. Multisensory brain mechanisms of bodily self-consciousness. Nature Reviews Neuroscience 13, 556–571. https://doi.org/10.1038/nrn3292

Blankertz, B., Lemm, S., Treder, M., Haufe, S., Müller, K.-R., 2011. Single-trial analysis and classification of ERP components--a tutorial. Neuroimage 56, 814–825. https://doi.org/10.1016/j.neuroimage.2010.06.048

Botvinick, M., Cohen, J., 1998. Rubber hands ‘feel’ touch that eyes see. Nature 391, 756– 756. https://doi.org/10.1038/35784

Boucsein, W., Fowles, D.C., Grimnes, S., Ben-Shakhar, G., roth, W.T., Dawson, M.E., Filion, D.L., Society for Psychophysiological Research Ad Hoc Committee on Electrodermal Measures, 2012. Publication recommendations for electrodermal measurements. Psychophysiology 49, 1017–1034. https://doi.org/10.1111/j.1469-8986.2012.01384.x

Braithwaite, JJ., Watson, DG., Jones, R., Rowe, M. (2013). A guide for analysing electrodermal activity (EDA) & skin conductance responses (SCRs) for psychological experiments. Psychophysiology, 49, 1017–1034.

Bröhl, F., Kayser, C., 2021. Delta/theta band EEG differentially tracks low and high frequency speech-derived envelopes. NeuroImage 233, 117958. https://doi.org/10.1016/j.neuroimage.2021.117958

Cao, Y., Summerfield, C., Park, H., Giordano, B.L., Kayser, C., 2019. Causal Inference in the Multisensory Brain. Neuron 102, 1076-1087.e8. https://doi.org/10.1016/j.neuron.2019.03.043

Cardini, F., Longo, M.R., Driver, J., Haggard, P., 2012. Rapid enhancement of touch from non-informative vision of the hand. Neuropsychologia 50, 1954–1960. https://doi.org/10.1016/j.neuropsychologia.2012.04.020

Cardini, F., Longo, M.R., Haggard, P., 2011. Vision of the Body Modulates Somatosensory Intracortical Inhibition. Cerebral Cortex 21, 2014–2022. https://doi.org/10.1093/cercor/bhq267

Cichy, R.M., Pantazis, D., 2017. Multivariate pattern analysis of MEG and EEG: A comparison of representational structure in time and space. NeuroImage 158, 441–454. https://doi.org/10.1016/j.neuroimage.2017.07.023

Cichy, R.M., Pantazis, D., Oliva, A., 2014. Resolving human object recognition in space and time. Nat Neurosci 17, 455–462. https://doi.org/10.1038/nn.3635

Costantini, M., Robinson, J., Migliorati, D., Donno, B., Ferri, F., Northoff, G., 2016. Temporal limits on rubber hand illusion reflect individuals’ temporal resolution in multisensory perception. Cognition 157, 39–48. https://doi.org/10.1016/j.cognition.2016.08.010

Critchley, H.D., Botan, V., Ward, J., 2021. Absence of reliable physiological signature of illusory body ownership revealed by fine-grained autonomic measurement during the rubber hand illusion. PLOS ONE 16, e0237282. https://doi.org/10.1371/journal.pone.0237282

Crucianelli, L., Krahé, C., Jenkinson, P.M., Fotopoulou, A.K., 2018. Interoceptive ingredients of body ownership: Affective touch and cardiac awareness in the rubber hand illusion. Cortex 104, 180–192. https://doi.org/10.1016/j.cortex.2017.04.018

D’Alonzo, M., Mioli, A., Formica, D., Di Pino, G., 2020. Modulation of Body Representation Impacts on Efferent Autonomic Activity. Journal of Cognitive Neuroscience 32, 1104–1116. https://doi.org/10.1162/jocn_a_01532

Ehrsson, H.H., 2004. That’s My Hand! Activity in Premotor Cortex Reflects Feeling of Ownership of a Limb. Science 305, 875–877. https://doi.org/10.1126/science.1097011

Ehrsson, H.H., Wiech, K., Weiskopf, N., Dolan, R.J., Passingham, R.E., 2007. Threatening a rubber hand that you feel is yours elicits a cortical anxiety response. PNAS 104, 9828–9833. https://doi.org/10.1073/pnas.0610011104

Eichert, N., Robinson, E.C., Bryant, K.L., Jbabdi, S., Jenkinson, M., Li, L., Krug, K., Watkins, K.E., Mars, R.B., 2020. Cross-species cortical alignment identifies different types of anatomical reorganization in the primate temporal lobe. eLife 9, e53232. https://doi.org/10.7554/eLife.53232

Evans, N., Blanke, O., 2013. Shared electrophysiology mechanisms of body ownership and motor imagery. NeuroImage 64, 216–228. https://doi.org/10.1016/j.neuroimage.2012.09.027

Faivre, N., Dönz, J., Scandola, M., Dhanis, H., Bello Ruiz, J., Bernasconi, F., Salomon, R., Blanke, O., 2017. Self-Grounded Vision: Hand Ownership Modulates Visual Location through Cortical β and γ Oscillations. J Neurosci 37, 11–22. https://doi.org/10.1523/JNEUROSCI.0563-16.2016

Fang, W., Li, J., Qi, G., Li, S., Sigman, M., Wang, L., 2019. Statistical inference of body representation in the macaque brain. PNAS 116, 20151–20157. https://doi.org/10.1073/pnas.1902334116

Ferri, F., Chiarelli, A.M., Merla, A., Gallese, V., Costantini, M., 2013. The body beyond the body: expectation of a sensory event is enough to induce ownership over a fake hand. Proc. R. Soc. B. 280, 20131140. https://doi.org/10.1098/rspb.2013.1140

Fuchs, X., Riemer, M., Diers, M., Flor, H., Trojan, J., 2016. Perceptual drifts of real and artificial limbs in the rubber hand illusion. Sci Rep 6, 24362. https://doi.org/10.1038/srep24362

Galigani, M., Ronga, I., Fossataro, C., Bruno, V., Castellani, N., Rossi Sebastiano, A., Forster, B., Garbarini, F., 2021. Like the back of my hand: Visual ERPs reveal a specific change detection mechanism for the bodily self. Cortex 134, 239–252. https://doi.org/10.1016/j.cortex.2020.10.014

Grabot, L., Kayser, C., 2020. Alpha Activity Reflects the Magnitude of an Individual Bias in Human Perception. J. Neurosci. 40, 3443–3454. https://doi.org/10.1523/JNEUROSCI.2359-19.2020

Graziano, M.S., Yap, G.S., Gross, C.G., 1994. Coding of visual space by premotor neurons. Science 266, 1054–1057. https://doi.org/10.1126/science.7973661

Graziano, M.S.A., Cooke, D.F., Taylor, C.S.R., 2000. Coding the Location of the Arm by Sight. Science 290, 1782–1786. https://doi.org/10.1126/science.290.5497.1782

Grootswagers, T., Cichy, R.M., Carlson, T.A., 2018. Finding decodable information that can be read out in behaviour. NeuroImage 179, 252–262. https://doi.org/10.1016/j.neuroimage.2018.06.022

Guggenmos, M., Sterzer, P., Cichy, R.M., 2018. Multivariate pattern analysis for MEG: A comparison of dissimilarity measures. NeuroImage 173, 434–447. https://doi.org/10.1016/j.neuroimage.2018.02.044

Guterstam, A., Björnsdotter, M., Gentile, G., Ehrsson, H.H., 2015. Posterior Cingulate Cortex Integrates the Senses of Self-Location and Body Ownership. Current Biology 25, 1416– 1425. https://doi.org/10.1016/j.cub.2015.03.059

Hipp, J.F., Siegel, M., 2013. Dissociating neuronal gamma-band activity from cranial and ocular muscle activity in EEG. Front. Hum. Neurosci. 7. https://doi.org/10.3389/fnhum.2013.00338

Holle, H., McLatchie, N., Maurer, S., Ward, J., 2011. Proprioceptive drift without illusions of ownership for rotated hands in the “rubber hand illusion” paradigm. Cognitive Neuroscience 2, 171–178. https://doi.org/10.1080/17588928.2011.603828ù

Holmes, N.P., Makin, T.R., Cadieux, M., Williams, C., Naish, K.R., Spence, C., Shore, D.I., 2012. Hand ownership and hand position in the rubber hand illusion are uncorrelated. Seeing and Perceiving 25, 52. https://doi.org/10.1163/187847612X646730

Horváth, Á., Ferentzi, E., Bogdány, T., Szolcsányi, T., Witthöft, M., Köteles, F., 2020. Proprioception but not cardiac interoception is related to the rubber hand illusion. Cortex 132, 361–373. https://doi.org/10.1016/j.cortex.2020.08.026

Kalckert, A., Ehrsson, H.H., 2014. The moving rubber hand illusion revisited: Comparing movements and visuotactile stimulation to induce illusory ownership. Consciousness and Cognition 26, 117–132. https://doi.org/10.1016/j.concog.2014.02.003

Kalckert, A., Ehrsson, H.H., 2012. Moving a Rubber Hand that Feels Like Your Own: A Dissociation of Ownership and Agency. Front. Hum. Neurosci. 6. https://doi.org/10.3389/fnhum.2012.00040

Kammers, M.P.M., Rose, K., Haggard, P., 2011. Feeling numb: Temperature, but not thermal pain, modulates feeling of body ownership. Neuropsychologia 49, 1316–1321. https://doi.org/10.1016/j.neuropsychologia.2011.02.039

Kanayama, N., Hara, M., Kimura, K., 2021. Virtual reality alters cortical oscillations related to visuo-tactile integration during rubber hand illusion. Scientific Reports 11, 1436. https://doi.org/10.1038/s41598-020-80807-y

Kanayama, N., Sato, A., Ohira, H., 2009. The role of gamma band oscillations and synchrony on rubber hand illusion and crossmodal integration. Brain and Cognition 69, 19– 29. https://doi.org/10.1016/j.bandc.2008.05.001

Kanayama, N., Sato, A., Ohira, H., 2007. Crossmodal effect with rubber hand illusion and gamma-band activity. Psychophysiology 44, 392–402. https://doi.org/10.1111/j.1469-8986.2007.00511.x

Keitel, A., Gross, J., Kayser, C., 2020. Shared and modality-specific brain regions that mediate auditory and visual word comprehension. eLife 9, e56972. https://doi.org/10.7554/eLife.56972

Kerby, DS. (2014). The simple difference formula: An approach to teaching nonparametric correlation. In Innovative Teaching.

Kilteni, K., Normand, J.-M., Sanchez-Vives, M.V., Slater, M., 2012. Extending Body Space in Immersive Virtual Reality: A Very Long Arm Illusion. PLOS ONE 7, e40867. https://doi.org/10.1371/journal.pone.0040867

Lenggenhager, B., Halje, P., Blanke, O., 2011. Alpha band oscillations correlate with illusory self-location induced by virtual reality. European Journal of Neuroscience 33, 1935–1943. https://doi.org/10.1111/j.1460-9568.2011.07647.x

Li, M., Wang, D., Ren, J., Langs, G., Stoecklein, S., Brennan, B.P., Lu, J., Chen, H., Liu, H., 2019. Performing group-level functional image analyses based on homologous functional regions mapped in individuals. PLoS Biol 17, e2007032. https://doi.org/10.1371/journal.pbio.2007032

Lloyd, D.M., 2007. Spatial limits on referred touch to an alien limb may reflect boundaries of visuo-tactile peripersonal space surrounding the hand. Brain and Cognition 64, 104–109. https://doi.org/10.1016/j.bandc.2006.09.013

Longo, M.R., Haggard, P., 2012. What Is It Like to Have a Body? Curr Dir Psychol Sci 21, 140–145. https://doi.org/10.1177/0963721411434982

Loomis, J.M., Philbeck, J.W., 1999. Is the anisotropy of perceived 3-D shape invariant across scale? Perception & Psychophysics 61, 397–402. https://doi.org/10.3758/BF03211961

Maris, E., Oostenveld, R., 2007. Nonparametric statistical testing of EEG- and MEG-data. J Neurosci Methods 164, 177–190. https://doi.org/10.1016/j.jneumeth.2007.03.024

McLaughlin, D.F., Kelly, E.F., 1993. Evoked potentials as indices of adaptation in the somatosensory system in humans: A review and prospectus. Brain Research Reviews 18, 151–206. https://doi.org/10.1016/0165-0173(93)90001-G

Nichols, T.E., Holmes, A.P., 2002. Nonparametric permutation tests for functional neuroimaging: a primer with examples. Hum Brain Mapp 15, 1–25. https://doi.org/10.1002/hbm.1058

O’Beirne, G.A., Patuzzi, R.B., 1999. Basic properties of the sound-evoked post-auricular muscle response (PAMR). Hearing Research 138, 115–132. https://doi.org/10.1016/S0378-5955(99)00159-8

Oostenveld, R., Fries, P., Maris, E., Schoffelen, J.-M., 2010. FieldTrip: Open Source Software for Advanced Analysis of MEG, EEG, and Invasive Electrophysiological Data. Computational Intelligence and Neuroscience 2011, e156869. https://doi.org/10.1155/2011/156869

Otsuru, N., Hashizume, A., Nakamura, D., Endo, Y., Inui, K., Kakigi, R., Yuge, L., 2014. Sensory incongruence leading to hand disownership modulates somatosensory cortical processing. Cortex 58, 1–8. https://doi.org/10.1016/j.cortex.2014.05.005

Park, H., Kayser, C., 2019. Shared neural underpinnings of multisensory integration and trial-by-trial perceptual recalibration in humans. eLife 8, e47001. https://doi.org/10.7554/eLife.47001

Parra, L.C., Spence, C.D., Gerson, A.D., Sajda, P., 2005. Recipes for the linear analysis of EEG. NeuroImage 28, 326–341. https://doi.org/10.1016/j.neuroimage.2005.05.032

Pavani, F., Spence, C., Driver, J., 2000. Visual Capture of Touch: Out-of-the-Body Experiences With Rubber Gloves. Psychol Sci 11, 353–359. https://doi.org/10.1111/1467-9280.00270

Peled, A., Pressman, A., Geva, A.B., Modai, I., 2003. Somatosensory evoked potentials during a rubber-hand illusion in schizophrenia. Schizophrenia Research 64, 157–163. https://doi.org/10.1016/S0920-9964(03)00057-4

Petkova, V.I., Björnsdotter, M., Gentile, G., Jonsson, T., Li, T.-Q., Ehrsson, H.H., 2011. From Part-to Whole-Body Ownership in the Multisensory Brain. Current Biology 21, 1118–1122. https://doi.org/10.1016/j.cub.2011.05.022

Press, C., Heyes, C., Haggard, P., Eimer, M., 2008. Visuotactile Learning and Body Representation: An ERP Study with Rubber Hands and Rubber Objects. Journal of Cognitive Neuroscience 20, 312–323. https://doi.org/10.1162/jocn.2008.20022

Preston, C., 2013. The role of distance from the body and distance from the real hand in ownership and disownership during the rubber hand illusion. Acta Psychologica 142, 177– 183. https://doi.org/10.1016/j.actpsy.2012.12.005

Pyasik, M., Ronga, I., Burin, D., Salatino, A., Sarasso, P., Garbarini, F., Ricci, R., Pia, L., 2021. I’m a believer: Illusory self-generated touch elicits sensory attenuation and somatosensory evoked potentials similar to the real self-touch. NeuroImage 229, 117727. https://doi.org/10.1016/j.neuroimage.2021.117727

Rao, I.S., Kayser, C., 2017. Neurophysiological Correlates of the Rubber Hand Illusion in Late Evoked and Alpha/Beta Band Activity. Front. Hum. Neurosci. 11. https://doi.org/10.3389/fnhum.2017.00377

Riemer, M., Bublatzky, F., Trojan, J., Alpers, G.W., 2015. Defensive activation during the rubber hand illusion: Ownership versus proprioceptive drift. Biol Psychol 109, 86–92. https://doi.org/10.1016/j.biopsycho.2015.04.011

Riemer, M., Trojan, J., Beauchamp, M., Fuchs, X., 2019. The rubber hand universe: On the impact of methodological differences in the rubber hand illusion. Neuroscience & Biobehavioral Reviews 104, 268–280. https://doi.org/10.1016/j.neubiorev.2019.07.008

Rohde, M., Di Luca, M., Ernst, M.O., 2011. The Rubber Hand Illusion: Feeling of Ownership and Proprioceptive Drift Do Not Go Hand in Hand. PLoS ONE 6, e21659. https://doi.org/10.1371/journal.pone.0021659

Rohe, T., Noppeney, U., 2015. Cortical Hierarchies Perform Bayesian Causal Inference in Multisensory Perception. PLOS Biology 13, e1002073. https://doi.org/10.1371/journal.pbio.1002073

Shibuya, S., Unenaka, S., Zama, T., Shimada, S., Ohki, Y., 2019. Sensorimotor and Posterior Brain Activations During the Observation of Illusory Embodied Fake Hand Movement. Front. Hum. Neurosci. 13. https://doi.org/10.3389/fnhum.2019.00367

Shimada, S., Fukuda, K., Hiraki, K., 2009. Rubber Hand Illusion under Delayed Visual Feedback. PLoS ONE 4, e6185. https://doi.org/10.1371/journal.pone.0006185

Sjouwerman, R., Lonsdorf, T.B., 2019. Latency of skin conductance responses across stimulus modalities. Psychophysiology 56, e13307. https://doi.org/10.1111/psyp.13307

Suzuki, K., Garfinkel, S.N., Critchley, H.D., Seth, A.K., 2013. Multisensory integration across exteroceptive and interoceptive domains modulates self-experience in the rubber-hand illusion. Neuropsychologia 51, 2909–2917. https://doi.org/10.1016/j.neuropsychologia.2013.08.014

Tieri, G., Tidoni, E., Pavone, E.F., Aglioti, S.M., 2015. Body visual discontinuity affects feeling of ownership and skin conductance responses. Scientific Reports 5, 17139. https://doi.org/10.1038/srep17139

Tsakiris, M., Haggard, P., 2005. The Rubber Hand Illusion Revisited: Visuotactile Integration and Self-Attribution. Journal of Experimental Psychology: Human Perception and Performance 31, 80–91. https://doi.org/10.1037/0096-1523.31.1.80

Valenzuela Moguillansky, C., O’Regan, J.K., Petitmengin, C., 2013. Exploring the subjective experience of the “rubber hand” illusion. Front. Hum. Neurosci. 7. https://doi.org/10.3389/fnhum.2013.00659

Van Horn, J.D., Grafton, S.T., Miller, M.B., 2008. Individual Variability in Brain Activity: A Nuisance or an Opportunity? Brain Imaging and Behavior 2, 327. https://doi.org/10.1007/s11682-008-9049-9

Zeller, D., Friston, K.J., Classen, J., 2016. Dynamic causal modeling of touch-evoked potentials in the rubber hand illusion. NeuroImage 138, 266–273. https://doi.org/10.1016/j.neuroimage.2016.05.065

Zeller, D., Litvak, V., Friston, K.J., Classen, J., 2015. Sensory Processing and the Rubber Hand Illusion—An Evoked Potentials Study. Journal of Cognitive Neuroscience 27, 573–582. https://doi.org/10.1162/jocn_a_00705

